# Blind spots in ecosystem services research and implementation

**DOI:** 10.1101/033498

**Authors:** Sven Lautenbach, Anne-Christine Mupepele, Carsten F. Dormann, Heera Lee, Stefan Schmidt, Samantha S.K. Scholte, Ralf Seppelt, Astrid J.A. van Teeffelen, Willem Verhagen, Martin Volk

## Abstract

Ecosystem service research has gained attraction, and the topic is also high on the policy agenda. Projects such as TEEB have generalized results of individual case studies to provide guidelines for policy makers and stakeholders. Seppelt et al. (2011) raised critical questions about four facets that characterize the holistic ideal of ecosystem services research: (i) biophysical realism of ecosystem data and models; (ii) consideration of trade-offs between ecosystem services; (iii) recognition of off-site effects; and (iv) comprehensive but shrewd involvement of stakeholders within assessment studies. An extended and updated analysis of ecosystem service case studies showed that the majority of these facets were still not addressed by the majority of case studies. Whilst most indicators did not improve within the span analyzed (1996–2013), we found a tendency for an increasing geographical spread of the case studies. Moreover, we incorporated an additional facet, namely the relevance and usability of case study results for the operationalization of the ecosystem service concept. Only a minority of studies addressed this facet sufficiently with no significant trend for improvement over time.

## Introduction

The goods and services that ecosystems provide for human well-being have attracted a lot of research in the last 15 years. A look at the available research databases shows that the growth of the literature on this topic is still accelerating (Chaudhary et al., 2015). Although the ecosystem services (ES) concept has been adopted in high-level policy frameworks, such as the Convention on Biological Diversity (https://www.cbd.int/), the Intergovernmental Platform on Biodiversity and Ecosystem Services (http://www.ipbes.net/) and the EU biodiversity strategy^1^, there is currently a mismatch between the considerable conceptual understanding of the ES concept in science, and the limited practical application thereof. This is reflected in the ongoing discussion on how the ES concept could be improved, mainstreamed and operationalized (Balmford et al., 2011; Bennett et al., 2015; Cowling et al., 2008; Daily et al., 2009; Fisher and Turner, 2008; Folke et al., 2011; Niemelä et al., 2010). This discussion has at least three different strands: (1) the *ES concept* itself, (2) the available *knowledge on ES* for practical implementation of the concept, and (3) *best practice for implementation* of the concept in practice. While the ES concept has become mainstream in environmental research and practice we still recognized a vivid scientific debate on the adequacy of knowledge and guidelines for implementations. Here we focus on the second point, the knowledge base needed for practical implementation of the concept (Ash et al., 2010).

Operationalization of knowledge from ES case studies is needed to support decision makers in managing limited resources provided by ecosystems in a sustainable way. Improving the knowledge base alone, does not automatically translate into an improved decision making (Jordan and Russel, 2014), but it is a necessary condition. Under the assumption that better informed decision makers come to better decisions, a sound knowledge base with respect to effects of global change (such as land-use, climate, demographic and socio-economic change) on the provisioning of ecosystem service is needed. Initiatives such as The Economics of Ecosystems and Biodiversity (TEEB, http://www.teebweb.org/) and the World Bank's Global Partnership for Wealth Accounting and the Valuation of Ecosystem Services (WAVES, https://www.wavespartnership.org/en) aim at providing such a knowledge base – relying as they do on the quality of results from individual case studies. Seppelt et al. (2011) raised concern about the soundness and usefulness of results of many ecosystem service studies. They aligned their criticism along four dimensions, biophysical realism, trade-offs, off-side effects and stakeholder involvement, and the limited consideration studies paid to each of these facets. In this study, we extended their approach by (1) analyzing changes since 2010 and (2) extending the analysis with respect to the relevance and usability of case study results for the operationalization of the ecosystem service concept. Our long-term perspective is to improve operationalization of ecosystem services, without compromising on the scientific knowledge base. Ecosystem service-based management requires more than a snapshot quantification. The biophysical quantification of an ES has to be dynamic (i.e. able to respond to changing environmental or technical conditions), and be supplemented by an analysis of synergies/trade-offs between ES, and consideration of export of negative decision effects (so-called off-site effects). Similarly, the social context of ES, the traditions, realizable management options, and governance structures must be taken into account, typically through elicitation and consultation with local stakeholders and decision-makers (Table 2).

## Methods and data

Seppelt et al. (2011) reviewed publications found through an ISI Web of Knowledge search of articles with the search phrase ‘ecosystem service’ OR ‘ecosystem services’ OR ‘ecosystem valuation’ in the title published up to 31.12.2010. We extended their database with a sample of articles selected by the same search phrases published between 01.01.2011 and 01.08.2013. The search phrase returned a total of 658 articles for the second period, from which we randomly selected 300 articles, of which again 118 were actual case studies. In total our analysis is therefore based on 271 case studies – 153 for the first and 118 for the second period. The original data set was updated for new attributes referring to relevance and usability, type of stakeholder involvement and trade-off analysis (see Table 1 for all attributes considered).

**Table 1.**
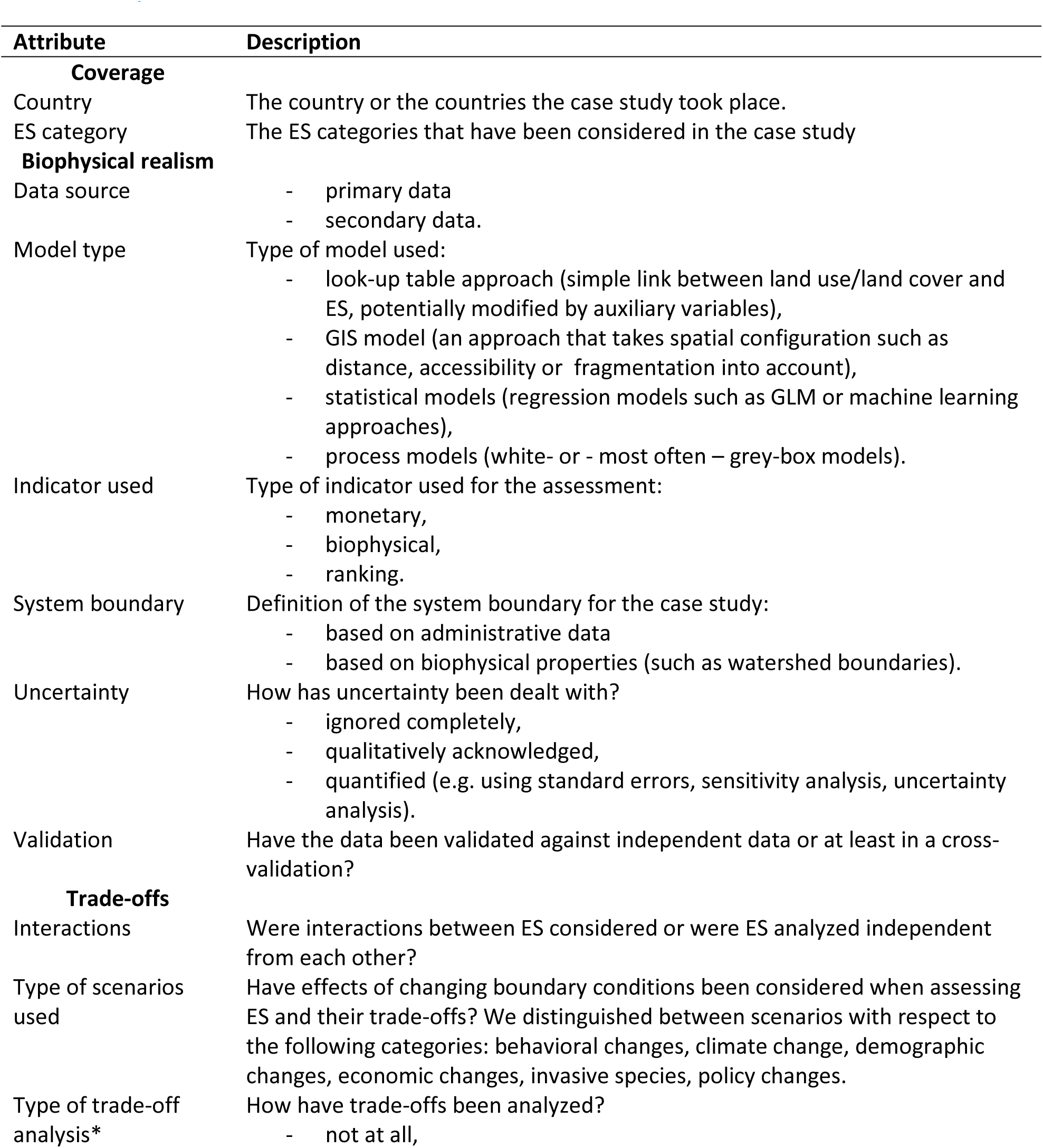
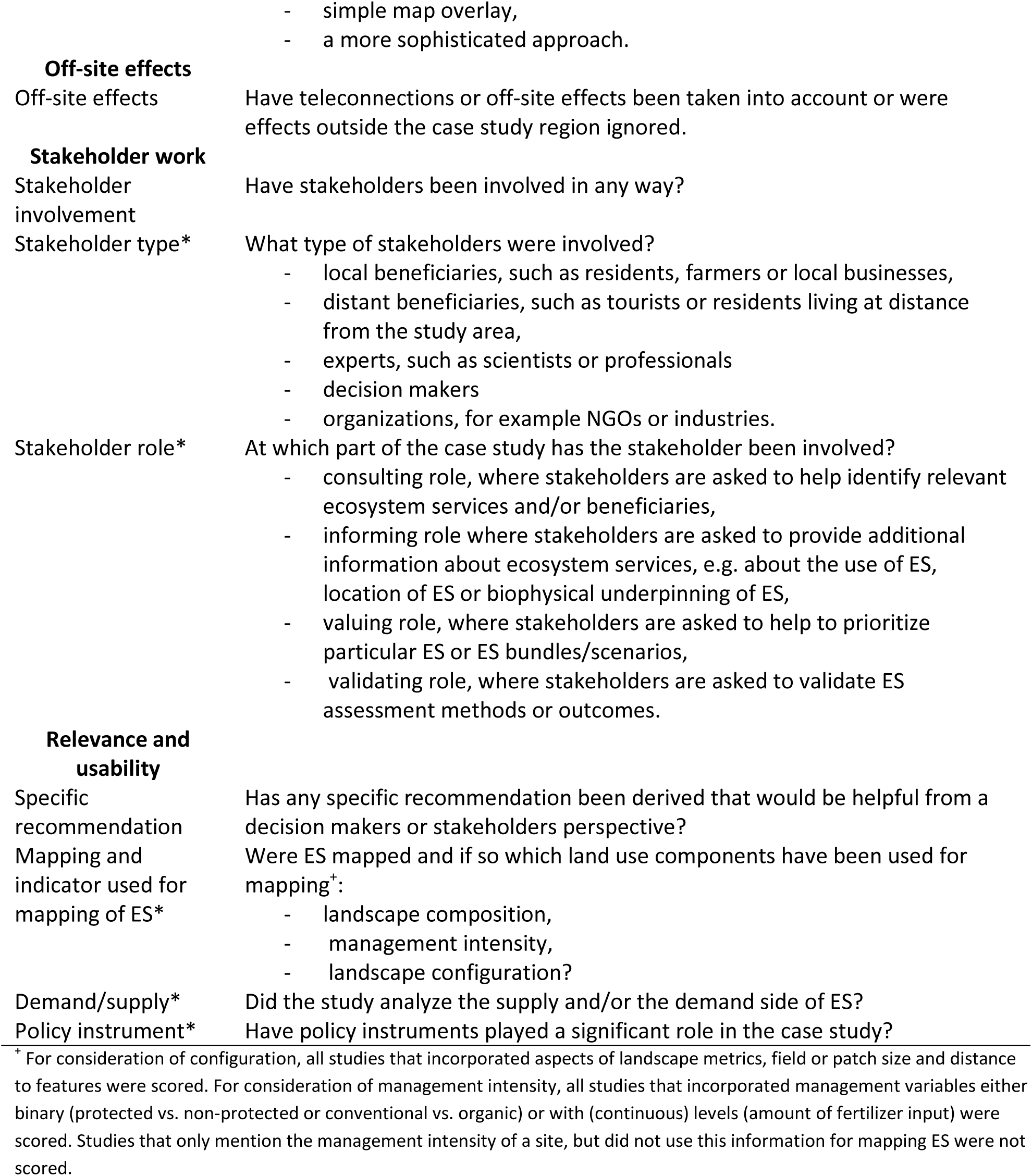
**Attributes used in the quantitative review. Additional attributes used in the review compared to** Seppelt et al. (2011) **are indicated by an asterisk**.

The data base is available as online supplementary material.

## Results and discussion

### Distribution of case studies over space and across ES categories

A global operationalization of ecosystem services requires case studies of similar quality to be conducted across the world. Most ecosystem services research was undertaken in the USA and China. This limits the ability of regional and local decision makers outside these two countries to identify most urgent threats since the knowledge base is relatively narrow in parts of the world. In recent years a tendency for a more even spread of ES studies across countries was observed (cf. supporting online material) – especially the US has lost its dominant role after 2004.

Beside geographical distribution, a representative range of ES need to be covered. There has been an unequal coverage of ES in research and this unequal distribution has not changed over the years. So far, research focused on ‘Food provisioning’, ‘climate regulation’ (mainly carbon sequestration), ‘biodiversity and nursery’ (mainly species richness indicators) and ‘opportunities for recreation and tourism’. Other ES such as ‘ornamental species’, ‘biochemical products and medicinal resources’ or ‘spiritual & artistic inspiration’ were rarely investigated. Possible reasons for this unequal distribution across categories are perceived importance of the ES category by researchers and/or stakeholders, different research traditions of study leaders, or financial, logistic and scientific challenges to quantify some ES in the field. A comprehensive ecosystem assessment is only possible when a representative set of ES is quantified, not only those that are particularly easy to estimate.

### Biophysical realism

Biophysical realism means that measurements, modeling and monitoring of ecosystem functions related to ecosystem services are close to the quantity measured and not abstract and unsubstantiated proxy indices (Seppelt et al., 2011). We used six indicators for biophysical realism: data source, model type, indicator used, system boundary, uncertainty and validation (Fig. 1, Table 2). Over time we observed little change in the biophysical realism of ES studies (cf. supporting online material). The majority of the case studies still used lookup tables together with simple land-cover classes as proxies, which further limits the reliability of a lookup table approach. Substantial shortcomings associated with the use of lookup tables have been reported by Konarska et al. (2002) and Eigenbrod et al. (2010a). Model approaches differ with respect to their ability to account for feedbacks (Liu et al., 2007), non-linear effects (Grêt-Regamey et al., 2014), spatial and temporal variability (Dale and Polasky, 2007; Eigenbrod et al., 2010) and to predict system behavior under changing boundary conditions.

**Figure 1.**
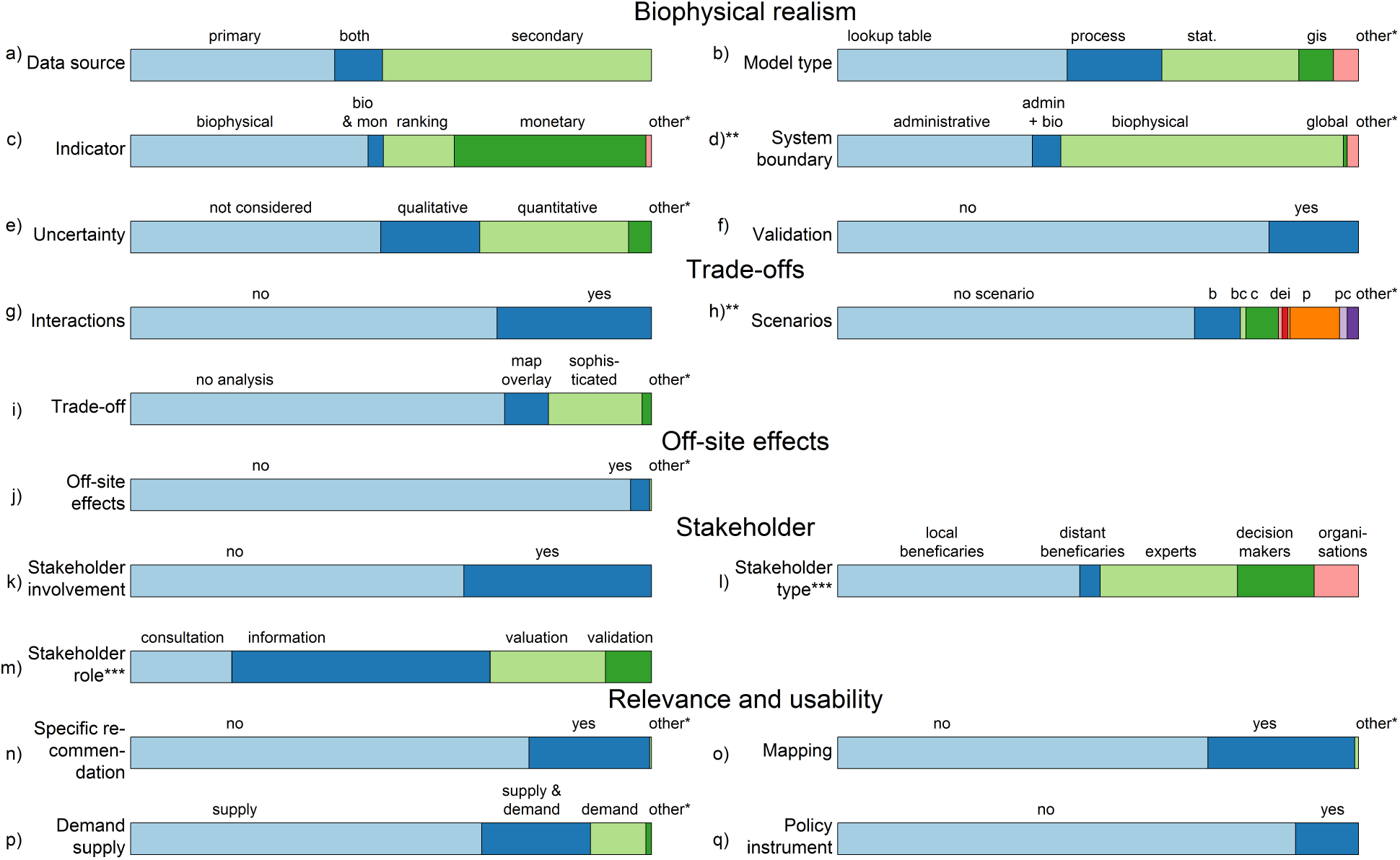
Percentage of the studies that belong to the specified factor level. The factor level ‘other’ refers to cases in which insufficient information to assign the article to a factor was given in the article. For scenarios the following types have been distinguished:: b -behavioral changes, c -climate change, d- demographic changes, e- economic changes, I – invasive species, p -policy changes, two letter combinations represent a combination of several scenario types in the same case study. *\*\** System boundary and scenarios belong not exclusively to one fact. *\*\*\** For stakeholder role and stakeholder type the percentage refers only to the number of studies that involved stakeholders.

The rest of the case studies relied on statistical models (26%), process models (18%) and GIS models (7%). There is no general best modeling approach – depending on the research question and the available data different approaches might be preferred. Given the complexity of human-environment interactions and the importance of feedbacks in ecosystem processes as well as in social processes, it has to be assumed that models that ignore such feedbacks are limited in their ability to correctly predict changes in both ES demand and supply.

Frequently, results obtained from ES models were neither validated nor are their uncertainties quantified. Assuming that all models are wrong but some are useful (Box, 1976) the usefulness of the models for the purpose of a case study needs to be demonstrated. The only way to estimate the reliability of any type of model is a test against independent data and an analysis of the uncertainty of model predictions (Bennett et al., 2012; Dormann et al., 2008; Hou et al., 2013; Jakeman et al., 2006; Kirchner et al., 1996; Laniak et al., 2013). Such a validation of results with independent data and an assessment of the attached uncertainty is a necessary prerequisite to judge conclusions drawn from model results. It should be mandatory to discuss results in the light of the quantified uncertainties and help thereby decision makers to decide if results are reliable enough to support the decision to be made.

### Trade-offs

Ecosystem services can be expected to interact (Bennett et al., 2009). This can be in form of trade-offs or synergies. The strength and even the direction of the interaction might change in space and time, and can be triggered by changing management strategies. In addition the interactions between ecosystem services can be non-linear and incorporate tipping points (Koch et al., 2009; Sabatier et al., 2013). Interactions between ES are of high relevance in multi-functional landscapes – ignoring them likely leads to suboptimal decisions. From a management perspective an ES case study should not only incorporate feedbacks between ES, but analyze the trade-offs or synergies between them as a crucial step for identifying promising management pathways(Bennett et al., 2009; Crouzat et al., 2014; Lee and Lautenbach, 2015).

Interactions between ES were largely ignored in ES case studies and this pattern did not change over time: the consideration of interactions between ES has been fluctuating around 30% with the exception of two peaks in 2003 and 2005 (cf. supporting online material). Even of the studies that looked at many different ES categories very few actually considered interactions between services. Considering interactions was particularly rare among case studies that used lookup tables, while it was more common in case studies that used statistical approaches or process models. In the case of process models the interactions were most frequently already built into the model. Our findings are in line with more specialized reviews such as that of Bommarco et al. (2013) on ecological intensification building on ES. They found nearly all studies had examined a single service process in isolation and that it had never been tested whether suites of below- and aboveground services contribute synergistically, or trade off, in their contribution to crop yield and quality.

Unexpectedly, studies that consider interactions between services and those that analyzed trade-offs were largely disconnected. This was mainly due to the use of simple map overlays in trade-off analysis: if two services compete for land (e.g. forestry and food production) they were identified as trading off. Even the more sophisticated approaches used to quantify trade-offs or synergies between ecosystem services had shortcomings: actual interactions of ecosystem services, such as the beneficial effects of forests on crop-pest control (Gagic et al., 2011) or pollination (Ricketts et al., 2008) and the positive effects of mosaics of land uses on biodiversity (Tscharntke et al., 2012b) were not considered, thus significantly underusing ecological system understanding (Raffaelli and White, 2013).

### Off-site effects

While the importance of off-site effects was already emphasized in Seppelt et al. (2011), it is still widely ignored in studies since 2011: only 10 studies (4%, 9 in 2011, 1 in 2012) incorporated off-site-effects. The decision to avoid local damage of ES might lead to a replacement of damaging activities to distant land systems. Taking this teleconnections or off-site effects into account, even in lump-sum approaches such as water or carbon footprints, might reveal underlying causes of developments and enable a better stakeholder selection as well as an improved system understanding (Lambin and Meyfroidt, 2011; Lapola et al., 2010; Meyfroidt and Lambin, 2009; Meyfroidt et al., 2013, 2010; Seto et al., 2012). Without consideration of such off-site effects there is substantial risk for the spatial spillover rebound effect (Maestre Andrés et al., 2012), in so far that policies intending to protect one type of biodiversity or ES in a certain area has even stronger negative impacts on such biodiversity or ES in another region. Emphasizing earlier comments, we believe that international agreements on ecosystem services need to be wise about the re-distribution of impacts in globally connected markets and that ES studies accordingly need to provide off-site analyses for their study systems.

### Stakeholder work

There is evidence that stakeholder participation can enhance the quality of environmental decisions by considering more comprehensive information inputs (Reed, 2008). Stakeholders were however only involved in 38% of the studies. The involvement of stakeholders was increasing up to 2005 and has remained relatively stable around 40% of the studies after 2005 (cf. supporting online material).

The type of stakeholders involved differed among studies. Overall, local beneficiaries were included the most (79% of studies), while distant beneficiaries were almost never included (5%). Experts were also often included (45%), while decision makers (25%) and organizations (15%) were addressed to a lesser extent. Most studies included only one type of stakeholders (47%), 39% included two types of stakeholders and 12% included three. Only two studies included four stakeholder categories. Stakeholders were more frequently involved in studies investigating the demand for ES than the supply side.

The role of stakeholders in the ES assessment varied between studies. Stakeholders were asked to prioritize ES (32%), identify ES (28%) or validate ES assessments (13%). Primarily stakeholders took an informing role in ES assessments (75%). This is not surprising as case study-specific information, as provided by local stakeholders, is often necessary for ES assessments. Local beneficiaries and decision makers were involved in all four stakeholder roles, while experts were more often approached for consultation and/or validation. Organizations and distant beneficiaries were never used for validation of ES assessments.

### Relevance and usability

If ecosystem assessments are to be of any use for decision making and management they need to be comprehensive. All relevant ES need to be quantified and valued, not only those that are particularly easy to estimate. Only then can the final step of operationalization be approached: incorporation of scientific results into decision making. Two aspects have to go hand in hand if ES assessments are a means to translate science into action: results of case studies should form a reliable basis that can be comprehended by decision makers; and decision makers should be included in form of stakeholders in the ES assessments. We analyzed several indicators to understand the implication of scientific results.

Mapping of ES is considered an important information tool for policy support (Maes et al., 2012a, 2012b; Verhagen et al., 2014). In total, only 28% of the studies mapped ES. Most of them relied on land-use composition (93%) largely ignoring management intensity (used by 37%) or landscape configuration (used by 29%). Just nine studies out of 70 included all three components of land use, of which six mapped only regulating services.

The ES concept links the provisioning of ES to societal needs supporting human well-being (Millenium Ecosystem Assessment, 2005). Goods and services provided by ecosystems have to reach their potential users otherwise the benefits cannot be realized. A complete ES case study should therefore aim at assessing both demand and supply for ES (Orenstein and Groner, 2014). While the supply side can be measured as biophysical indicators, social demand for ES can be valued using economic valuation techniques in real or hypothetical markets (Bateman et al., 2011; Turner et al., 2010), or based on non-monetary indicators such as the assessment of people's perceptions of the importance of different services (Martín-López et al., 2012; Scholte et al., 2015).

Most studies were found to ignore either ES demand or ES supply. The majority of ES case studies (70%) focused on the supply side of ES, while 12% studied only the demand side of ES, and only 18% looked at aspects of both sides at the same time. Over time, the share of studies on ES supply steadily increased, while the share of studies on both ES supply and ES demand steadily decreased. The share of ES demand studies showed an initial increase till 2007, but afterwards slightly decreased. Following Wolff et al. (2015), it seems we need to improve the understanding of the demand for ecosystem services.

The majority of studies (69%) did not consider any type of scenario but focused instead on an analysis of the current state. ES assessments were treated mainly as a static analysis without considering changes on both the demand and the supply side of ES. The use of scenarios has been widely fluctuating between 60% and 4% (cf. supporting online material). Land systems are dynamic by nature and change over time. Similar, demand and supply for ES are likely to change with time. A sustainable management of land systems has therefore to consider potential future changes of the system (McCauley, 2006). Changes in the demand and/or the supply side of ES have to be expected due to various reasons, such as climate change (e.g. Prather et al., 2013), demographic changes (shrinking/growing population, aging of population), behavioral changes (e.g. changes in consumption behavior: Tscharntke et al. 2012), economic development (e.g. the financial crisis and its effects on resource availability for environmental conservation), or policy changes (e.g. bioenergy: Banse et al., 2008; Campbell and Doswald, 2009; Landis et al., 2008; Lautenbach et al., 2013).

Scientific results may translate into specific recommendations on best practice. Few studies did provide any kind of specific recommendation (33%). After a strong peak in 2003 the percentage of studies that provided specific recommendations has been fluctuating without a clear trend (cf. supporting online material). This is in line with the results from Laurans et al. (2013), who looked specifically at ecosystem service valuation studies and found that it is common to present an economic valuation and recommend its use without this use being either explicated or contextualized, and concrete examples being neither provided nor analyzed. Notably, demand-side studies were *not* more likely to provide specific recommendations. However, studies involving stakeholders *were* more likely to provide specific recommendations (31%) than studies that did not involve stakeholders (17%). Studies that mapped ES did not provide specific recommendations significantly more frequent than others. This lack of specific recommendations from mapping studies contrast with the emphasis placed on mapping as an information tool for policy support (e.g. Maes et al., 2012a, 2012b).

The majority of the case studies (87%) did not consider any policy instrument. Only the relatively broad category of payments for ecosystem services (PES) was mentioned repeatedly (12 times), while all other instruments were mentioned maximally once. The consideration of policy instruments in ES studies seems to have slightly increased from 2005 onwards and has been fluctuating between 15% and 20% (cf. supporting online material).Management decisions are often concerned with the question of how to reach a specific objective, i.e. through which policy instrument. ES can be thought of as important dimensions to compare the (assumed) outcomes of policy instruments from the perspective of society. Studies that assess the effects of policy instruments on ES provide highly relevant information for decision makers and stakeholders. The lack of studies on specific policy instrument clearly hamper transferring the ES concept from science to policy and practice.

**Table 2.**
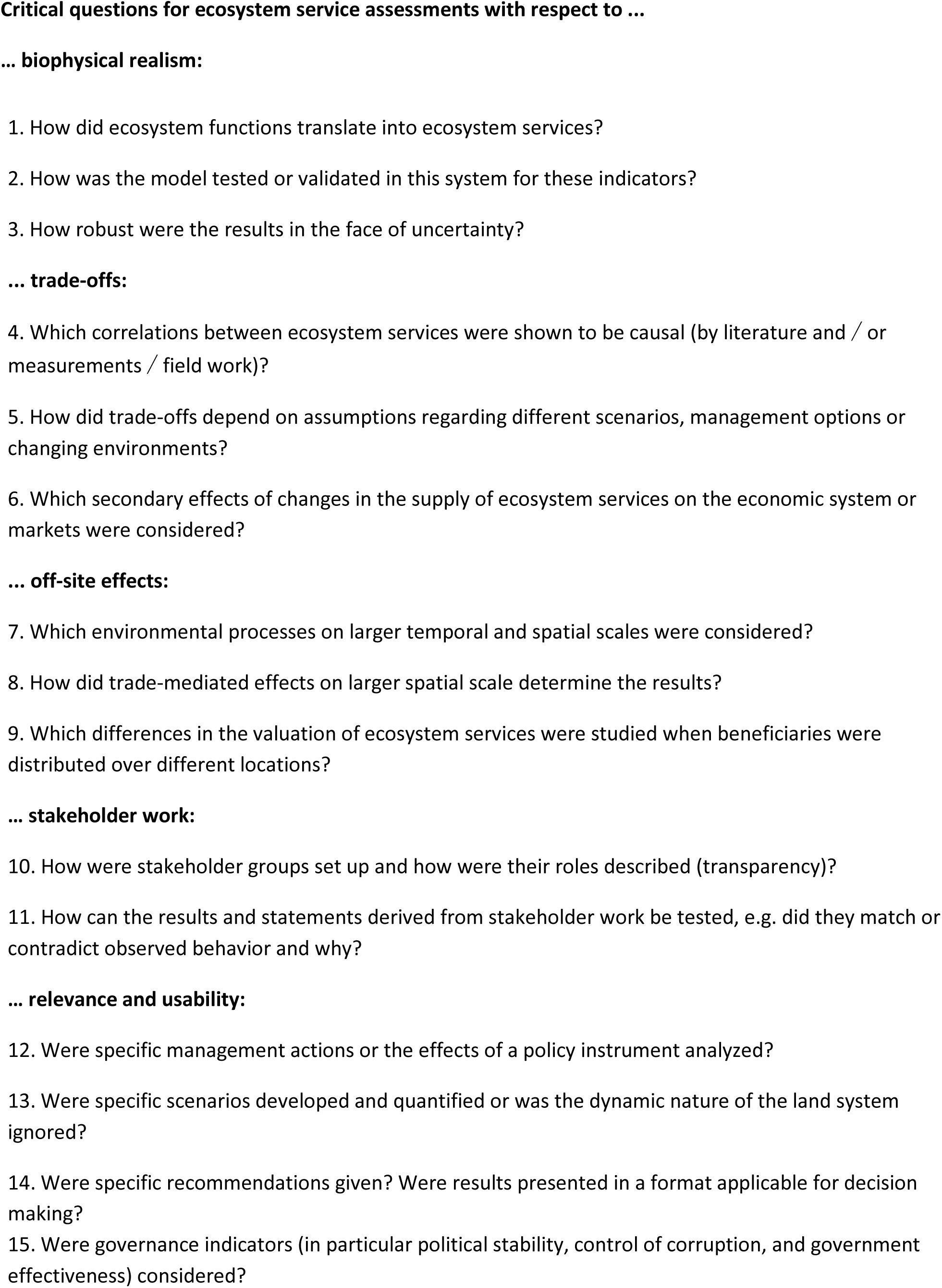
**Critical questions for reviewing ecosystem service assessments studies at the regional scale, based on** (Seppelt et al., 2011)

## Conclusions

The number of published ES studies has continued to rise over the last years. Unfortunately, shortcomings with respect to biophysical realism, trade-off analysis, off-site effects, stakeholder involvement as well as relevance and usability still persist. There is no support for the hypothesis that more recent studies overcome these shortcomings. In addition, the uneven coverage of different land systems and of ES categories hinders the efficient steering of resources for the conservation of ES. One has to be careful to assess the current state of knowledge on ES – it is not enough to look at the number of studies on specific ES categories or to look at the number of studies in a region. Instead, it is important to filter out the case studies that fulfill specific requirements. To operationalize the concept of ES the mentioned blind spots need to be addressed by upcoming studies. The list of critical questions provided (Table 2) hopefully helps to raise the awareness of the blind spots.

## Acknowledgements

The work has been funded by the EU-FP7 project OPERAs under grant agreement number FP7-ENV-2012–3 08393.

## Footnotes

1 http://ec.europa.eu/environment/nature/biodiversity/comm2006/2020.htm

